## Supplementary Materials for "Mechanism of connexin channel inhibition by mefloquine and 2-aminoethoxydiphenyl borate"

### Materials and Methods

**Expression plasmids.** Human connexin-32 (Cx32, Uniprot ID P08034) or human connexin-43 (Cx43, Uniprot ID P17302), fused to a C-terminal YFP and twinStrep tag, were cloned into a pACMV plasmid. The IRES (internal ribosomal entry site) containing construct was generated by subcloning the tag-free connexin coding sequence using NheI and MreI restriction sites prior to the IRES sequence followed by YFP sequence in pACMV vector. All mutations were generated by standard PCR-based procedures in the IRES-containing connexin constructs, using KAPA HiFi Hotstart Ready Mix (Roche), DpnI (NEB, #R0176S).

**Cell culture.** For protein expression, the adherent cultures of HEK293F cells were maintained in 15 cm cell culture plates in Dulbecco's Modified Eagle's Medium (DMEM, BioConcept), supplemented with 10 % Fetal Calf Serum (FCS) and 1 % Penicillin-Streptomycin (PenStrep, PAN Biotech) at 37 °C and 5 % $CO_2$. Prior to transfection, the medium in 100 plates of adherent HEK293F cultures was exchanged to DMEM supplemented with 2 % FCS and 1 % PenStrep. The cells were transfected using branched polyethylene imine (PEI, Sigma-Aldrich), by preparing a DNA dilution using 50 µg of YFP-twinStrep construct DNA per plate in non-supplemented DMEM, and a PEI dilution with 100 µg PEI (DNA: PEI = 1:2 [w/w]) in non-supplemented DMEM. After mixing and incubating the DNA- and PEI-containing solutions for 5 minutes to allow DNA-PEI complex formation, the mixture was added to the plates in a drop-wise manner. The cells were incubated at 37 °C and 5 % $CO_2$ for 48 h, harvested by scraping and centrifugation (4 °C, 500xg). The cell pellets were frozen and stored at – 80 °C until the day of experiment.

For immunocytochemistry (ICC) experiments the cells were seeded on glass coverslips in wells of a 6-well plate at a density of 0.5 million cells per well in DMEM supplemented with 10 % FCS and 1 % PenStrep. For transfection, the medium was exchanged to DMEM supplemented with 2 % FCS and 1 % PenStrep and transfected as described, using 3 µg of IRES construct DNA per well, and placed at 37 °C and 5 % $CO_2$ for 36 h of protein expression.

For functional experiments, namely hemichannel and gap-FRAP (fluorescence recovery after photobleaching) assays, the cells were seeded into poly-L-lysine pre-coated ibidi µ-dishes (#80136-IBI) or µ-slides (#80826-90), using 0.04 or 0.2 million cells per well or dish respectively, in DMEM supplemented with 10 % FCS 1 % PenStrep. The cells were transfected using PEI as described above, using 0.5 µg or 2 µg of IRES construct DNA per well or dish, respectively. The cells were incubated at 37 °C and 5 % $CO_2$ for 36 h of prior to the experiments.

**Protein purification.** The cell pellets (corresponding to a total of 100 plates of adherent cell culture) were resuspended in 100 ml buffer A (25 mM Tris-HCl, 150 mM NaCl) supplemented with a tablet of EDTA-free Complete protease inhibitor cocktail (04693132001; Roche, Basel, Switzerland). The cells were lysed with Vibra-Cell sonicator, using a 0.5 s pulse per plate, with a 0.5 s break in-between, at a 35 % amplitude. The lysate was clarified by ultracentrifugation at 82 000 x g and 4 °C for 50 min (Beckman Coulter Ti45 rotor). The supernatant was discarded, and the pellet (membrane fraction) resuspended in buffer A, supplemented with the Complete protease inhibitor cocktail, using ultraTurax homogenizer. The membranes were solubilized using a mixture of 1 % dodecyl-β-D-maltopyranoside (DDM) and 0.2 % cholesteryl hemisuccinate (CHS) (final concentration) for 1 h at 4 °C with constant rotation. The solubilized membranes were subjected to another round of ultracentrifugation and the supernatant was incubated with 1 ml of anti-GFP nanobody-coupled CNBr Sepharose resin [1] for 30 min at 4 °C with constant rotation. The resin was collected in a gravity column and washed with 40 column volumes of buffer B (25 mM Tris-HCl pH 8.0, 150 mM NaCl, 0.1 % digitonin). The protein was eluted by overnight cleavage with HRV 3C (200 µg) at 4 °C. The eluted protein was concentrated using a 100 kDa cut-off AmiconUltra concentrator and purified by size exclusion chromatography (SEC) using a Superose 6 Increase 6/300 GL (GE Healthcare), pre-equilibrated with buffer B. The fractions, corresponding to the protein peak, were pooled together, concentrated using a 100 kDa cut-off Amicon Ultra concentrator and immediately used for further experiments.

**Cryo-EM sample preparation and data collection.** The purified Cx32 in digitonin was concentrated to a concentration of 5 mg/ml; the purified Cx43 in digitonin was concentrated to 2 mg/ml. Mefloquine (MFQ) and 2-aminoethoxydiphenyl borate (2APB) were added to separate samples at a final concentration of 1 mM and incubated with the protein on ice for 30 min. A 3.5 µl aliquot of the protein was applied to a glow discharged Quantifoil R1.2/1.3 200-mesh grid and plunge frozen in liquid ethane using Vitrobot Mark IV (Thermo Fisher Scientific; 3 s blot time and blot force 20). The cryo-EM datasets were all collected using Titan Krios electron microscope equipped with a K3 direct electron detector (Gatan) and a GIF-Quantum energy filter (slit width 20 eV) at ScopeM, ETH Zürich. The range of defocus was set from -0.5 µm to -2.5 µm and the data collected in super-resolution mode using EPU 2.0, with the movies dose fractioned into 40 frames. The dose for the two collected Cx32-MFQ datasets were 50 e/Å$^2$ and 48 e/Å$^2$, for three Cx32-2APB datasets 52 e/Å$^2$, 44 e/Å$^2$ and 48 e/Å$^2$, and for two Cx43-MFQ datasets 60 e/Å$^2$.

**Cryo-EM data processing.** The images were separated into different optics groups according to the EPU beam shift values with a script developed by Dr. Pavel Afanasyev (CryoEM Tools v0.0.1) [2]. The movies were corrected using MotionCor2 [3] and CTFFIND4.1 [4] or Gctf [5] were used for CTF estimation for Cx32-MFQ and 2APB datasets, respectively, and CTFFIND4.1 for Cx43-MFQ datasets. In the case of Cx32 datasets approximately 1000 particles (MFQ: 835, 2APB: 1035) were manually picked in Relion4.0 [6] and subjected to 2D classification, where the 2D classes showed clear features of GJCs and HCs. In case of Cx43 + MFQ datasets, Laplacian-of-Gaussian autopicking was performed to obtain 20849 particles, which were subjected to 2D classification. For all datasets best classes were selected for reference-based autopicking in Relion 4.0. After several rounds of 2D classification, the remaining particles from the two Cx32-MFQ datasets, three Cx32-2APB datasets, or two Cx43-MFQ datasets were merged. Following several additional 2D classification jobs, 2D classes with clearly distinguished GJC and/or HC features were selected and extracted for 3D classification separately. The Cx32-apo GJC and HC maps [7], low pass filtered to 40 Å, were used to 3D classify GJC and HC particles in Cx32 datasets separately in Relion 4.0, with imposed D6 or C6 symmetry, respectively. The Cx43-apo GJC map in digitonin [8], low pass filtered to 40 Å, was used to classify the Cx43 GJC particles of the Cx43-MFQ dataset. The best 3D classes were selected for masked 3D refinement with imposed symmetry, followed by CTF refinement and Bayesian polishing in Relion 4.0. ResMap [9] was used for obtaining local resolution maps.

The final refined Cx32 maps with imposed symmetry were also used for protomer focused classification. The particles were symmetry expanded using ‘relion_particle_symmetry_expand’ command. The reference map of the single subunit was generated based on the model in Chimera [10], using ‘color zone’ and ‘split map’ functions. The reference was used for generation of the mask, which was low pass filtered to 10 Å, with addition of 5 pixels and a soft edge of 20 pixels. The symmetry expanded particles were subtracted, re-centred based on the mask, and downscaled to 200 x 200 pixels. The subtracted particles were subjected to 3D classification without alignment and symmetry imposition, classifying to 4, 6 or 8 classes.

Details of the image processing pipeline are shown in Extended Data Figs. 5-9, 12, 14-15, and Extended Data Table 1. Data visualization was performed using PyMOL [11] and UCSF ChimeraX [12].

**Model building and refinement.** The models of Cx32-apo GJC (PDB ID 7ZXM) and HC (PDB ID 7ZXN) were used as templates for building the Cx32-MFQ or Cx32-2APB GJC and HC models, respectively, in COOT [13]. The cytoplasmic regions (GJC: M1-R15, H100-S128 and A218-C283; HC: E109-K121 and A221-C283) were not built as they were not resolved in the cryo-EM map. The (-)-(11*S*,12*R*) MFQ enantiomer (chemical ID YMZ) was fit into the density based on Cx36 GJC structure in complex with MFQ (PDB ID: 8QOJ) [14]. 2APB (chemical ID F4Z) was fit in two densities, not present in Cx32-apo GJC and HC maps by rigid body fit followed by real space refinement. Cx43-MFQ GJC model was built based on Cx43-apo GJC structure in detergent as a template (PDB ID: 7Z1T). In Cx43-MFQ GJC model we did not build the cytoplasmic regions as they were not resolved in the reconstruction (K105-G150 and G235-I382). (-)-(11*S*,12*R*) MFQ was placed into the density based on the MFQ position in the Cx36-MFQ GJC model (PDB ID: 8QOJ). Any outstanding clashes which were not resolved in PHENIX were resolved using Chiron [15]. All structures were refined using 'phenix.real_space_refine' and evaluated with MolProbity in PHENIX [16,17].

**Electrostatic surface potential calculations**. The molecules were prepared for electrostatic calculations using PDB2PQR [18] with AMBER ff99 force field [19]. Calculation of the electrostatic surface potentials of protein in complex with ligands was performed using APBS Tools 2.1 [20] in PyMOL with the nonlinear Poisson-Boltzmann Equation.

**Intrinsic tryptophan fluorescence quenching-based binding assays.** The protein was diluted to a final concentration of 0.2 mg/ml in buffer B, centrifuged at 24 000 x g in benchtop centrifuge for 5 min, and transferred to a quartz glass cuvette. The spectra from 300 nm to 500 nm were collected using Carry Eclipse fluorescence spectrophotometer at an excitation wavelength of 295 nm. MFQ and 2APB were titrated by addition of small volumes (no bigger than 0.3 µl at time) from stock dilutions, with final concentration of the individual drug ranging from 61.5 nM to 96.7 µM. All data was blank corrected, and the values at 340 nm used for calculation of fluorescence quenching. For 2APB-related data, fluorescence emission shift was additionally calculated using the following equation:

$$Fluorescence\ emission\ shift = \frac{Iem(340\ nm) + Iem(350\ nm)}{Iem(330\ nm)}$$

with Iem, denoting the fluorescence emission intensity at the respective wavelength. The data were analyzed in GraphPad Prism 9.0.0 (for iOS, GraphPad Software, San Diego, California USA, www.graphpad.com) using a one-site specific binging model for MFQ and two-site specific binding model for 2APB. The decision on the model was done by analyzing the statistics of the fit as well as the fraction of molecules, occupying the site.

**Immunocytochemistry.** The cells were washed with PBS and fixed on glass coverslips with 4 % PFA in PBS (pH 7.4). The coverslips were placed in 100 mM Tris pH 9.0, 5 % [w/v] urea at 95 °C for antigen retrieval. The coverslips were washed three times using 50 ml PBS and placed in 100 µM digitonin PBS for membrane permeabilization for 10 min. The coverslips were washed three times using 50 ml PBS, placed in 1 % BSA, 22.25 mg/ml glycine in PBS for 30 min, and incubated with rabbit anti-connexin antibody (Cx32: #34-5700, 1:250 dilution, Invitrogen) in 1 % BSA in PBS overnight at 4 °C in a humidity chamber. The coverslips were washed again three times with 50 ml PBS and incubated for 1 h with Alexa Fluor® 647-conjugated goat-anti rabbit secondary antibody (#ab150079, Abcam, 1:2000 dilution) and again rinsed three times with PBS. The coverslips were stained with 50 µg/ml Hoechst 33342 (Sigma) in PBS, washed three times with PBS and mounted on glass slides with Fluoromount-G mounting medium (Invitrogen). The samples were stored at 4 °C prior to imaging.

The z-stacks were collected with Leica Stellaris 5 confocal microscope using LAS X data acquisition software (4.2.1.23819 – build 23180), a HC PL APO 63X/1.4 oil CS2 objective and a HyD detector. The stacks were collected in a frame-sequential data acquisition scheme every 0.3 µm for 7 µm, with the center of the stack defined and the z height with highest Hoechst 33342 signal, using 405 nm (Hoechst 33342), 488 nm (EYFP) and 638 nm (Alexa 647) laser lines, and a pinhole airy of 1.06 AU (101.4 µm), pixel dwell time of 3.1625 µs and a scan speed of 400 Hz. The stacks were imported to Fiji (ImageJ)[21] and different channels of the same stack were merged. GJ plaques (connexin-stained lines which appeared in the or more z-slices) were measured and counted. The measurements were imported into GraphPad Prism 9.0.0 and different conditions were compared using one-way ANOVA followed by Tukey's multiple comparisons test for GJ plaque number and Dunn's multiple comparisons test for GJ plaque length.

**Hemichannel activity assays.** The cells grown in ibidi-slides were washed twice with 200 µl of PBS-E (PBS pH 7.4, 5 mM EGTA) per well and incubated in PBS-E for 5 min. PBS-E was replaced with PBS-E containing 50 µg/ml sulforhodamine 101 (SR101) and 5 µg/ml Hoechst33342 for 10 min, after which the cells were washed four times with PBS (pH 7.4) and imaged in FluroBright™ DMEM supplemented with 1% FCS and 1 % PenStrep. In case of MFQ treatment, MFQ in DMSO was added to PBS-E/PBS-E-SR101 at a 30 µM final concentration. 2APB in DMSO was added at a final concentration of 100 µM. For each condition 10 micrographs were collected using Leica Stellaris 5 confocal microscope with the same configuration as described above, using 405 nm (Hoechst 33342), 488 nm (EYFP) and 561 nm (SR101) laser lines and the same imaging parameters. The images were processed and measured using an in-house developed script (https://github.com/lavrihap/hc-data-processing.git) [7] and the results were analyzed using GraphPad Prism 9.0.0. Three independent experiments were conducted for each condition. After confirmation that independent experiments yield the same result, the replicate measurements were merged together into one dataset. The significance of the differences in dye-uptake between the individual conditions was assessed using one-way ANOVA followed by Games-Howell's multiple comparisons test.

**Gap-FRAP assays.** The cells were treated as described for the hemichannel assays. In case of MFQ treatment, MFQ was added to FluroBright™ DMEM at a final concentration of 30 µM, and 2APB at 100 µM, and the cells were incubated for 5 min before start of data collection. Neighboring cells in direct contact, containing YFP fluorescence, were considered to be connected by GJCs. One of the cells forming close contacts was selected for bleaching of the SR101 dye. Six micrographs were collected at a low laser intensity, followed by bleaching of SR101 in the selected cell at 100 % 561 nm laser line intensity in 5.16 s intervals. The fluorescence recovery was recorded at the initial lower laser line intensity every 20.16 s for 45 frames. FRAP time series were imported into Fiji (ImageJ) and measured in three regions: bleached cell (ROI1), neighboring YFP-containing cells (ROI2) and background (ROI3). The fluorescence recovery was calculated using EasyFRAP-web software [22] using full-scale normalization. The normalized curves were exported to GraphPad Prism 9.0.0 and a two-phase or one-phase association curve (whichever statistically described the data better) was fit to the data. The fits for individual conditions were compared using extra sum-of-squares F-test. The plateaus of FRAP recovery for all conditions were compared using one-way ANOVA followed by Tukey's multiple comparisons test.

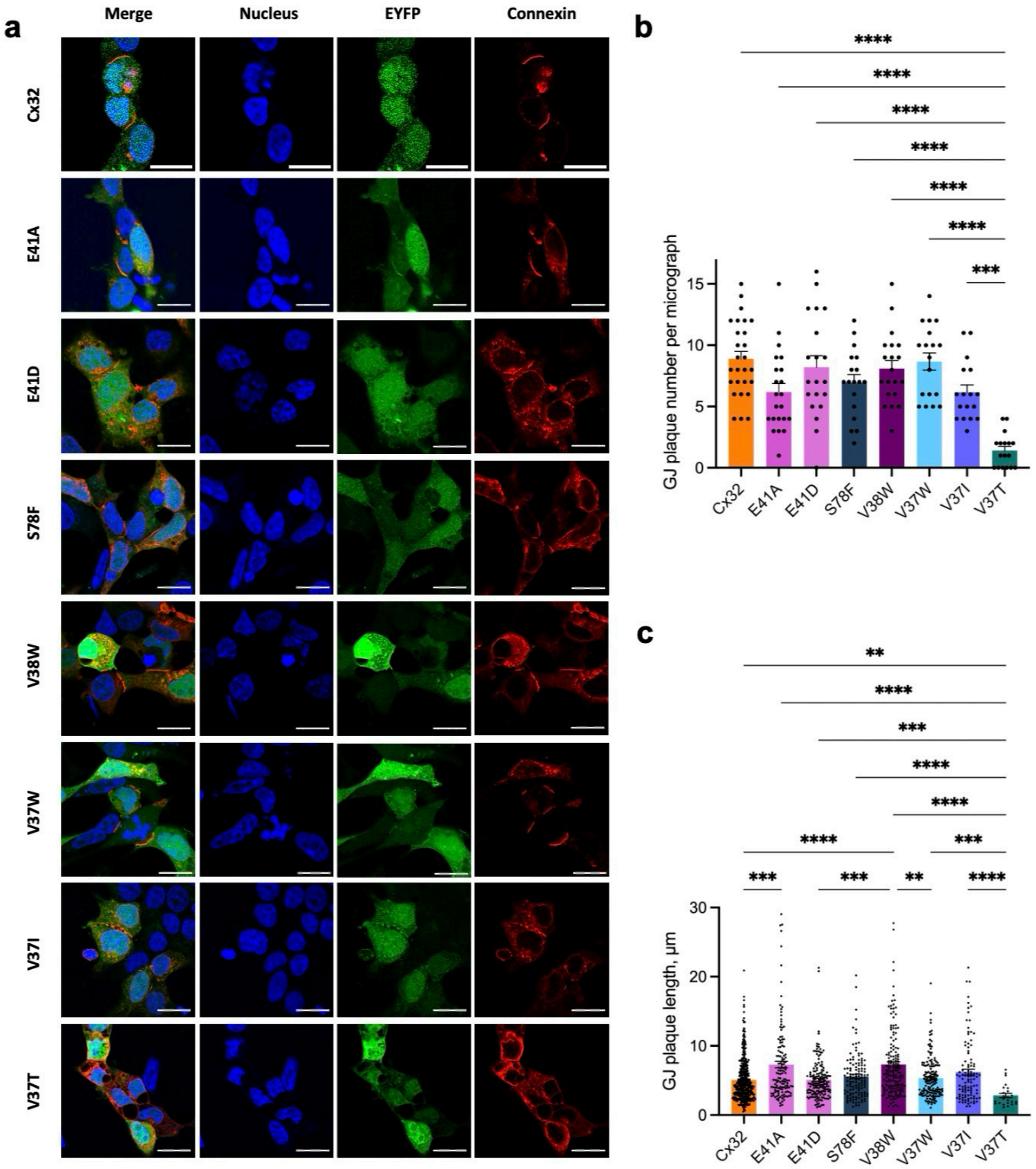


**Extended Data Fig. 1. Immunocytochemistry of Cx32 and site M mutants in HEK293F cells. a,** Representative immunocytochemistry micrographs of wild-type Cx32 and site M mutants, expressed in HEK293F cells. The cells were stained with Hoechst 33342 (blue – nuclei), and rabbit primary anti-connexin 32 antibody, followed by staining with Alexa-647 conjugated goat anti-rabbit secondary antibody (red – connexin). The cells, expressing Cx32, co-express YFP (green), encoded in the same vector after the IRES. Scale bar = 18 µm. **b,** Number of GJ plaques per micrograph for different constructs: Cx32, n = 27; E41A, n = 22; E41D, n = 20; S79F, n = 18; V38W, n = 20; V37W, n = 18; V37I, n = 17; V37T, n = 17. **c,** Length of GJ plaques in Cx32 and site M mutants: Cx32, n = 396; E41A, n = 136; E41D, n = 164; S79F, n = 125; V38W, n = 162; V37W, n = 159; V37I, n = 105; V37T, n = 24. Significance was determined using one-way ANOVA and followed by Tukey's (b) and Dunn's (c) multiple comparisons test: **** - $P < 0.0001$, *** - $P < 0.001$, ** - $P < 0.01$.

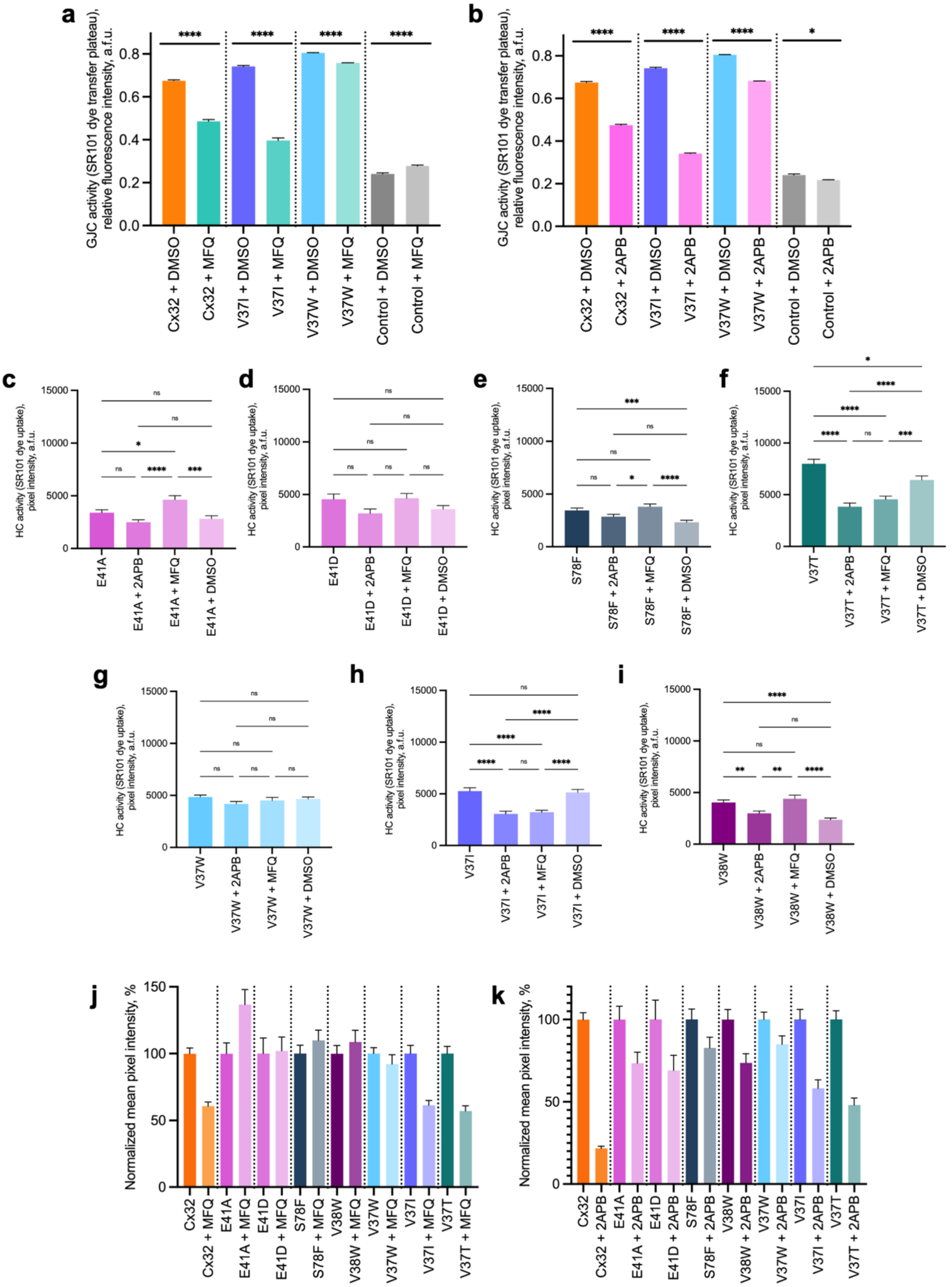


**Extended Data Fig. 2. Effects of MFQ and 2APB on the GJC and HC activity of the wild-type Cx32 and site M mutants. a,** Comparison of gap-FRAP data plateau values upon treatment with 30 μM MFQ (n = 41). **b,** Comparison of gap-FRAP data plateau values upon treatment with 100 μM 2APB (n = 41). Gap-FRAP values were compared using one-way ANOVA followed by Tukey's multiple comparisons test: **** - P < 0.0001, * - P < 0.05. **c-j,** HC activity of E41A (c, no treatment: n = 605, 2APB: n = 629, MFQ: n = 593, DMSO: n =

597), E41D (d, no treatment: n = 355, 2APB: n = 368, MFQ: n = 336, DMSO: n = 442), S79F (e, no treatment: n = 916, 2APB: n = 817, MFQ: n = 818, DMSO: n = 890), V38W (f, no treatment: n = 644, 2APB: n = 685, MFQ: n = 435, DMSO: n = 618), V37W (g, no treatment: n = 1042, 2APB: n = 1017, MFQ: n = 576, DMSO: n = 1279), V37I (h, no treatment: n = 873, 2APB: n = 708, MFQ: n = 768, DMSO: n = 852), V37T (i, no treatment: n = 701, 2APB: n = 615, MFQ: n = 713, DMSO: n = 767). *n – number of measured cells; all experiments were performed in experimental triplicates.* The statistical significance of differences between datasets was evaluated using Games-Howell's multiple comparisons test: **** - P < 0.0001, *** - P < 0.001, ** - P < 0.01, * - P < 0.05 , ns - P > 0.05. **j-k,** A histogram plot of HC activity data of Cx32 and Cx32 site M-mutants upon MFQ (j) or 2APB (k) treatment, normalized to HC dye uptake without treatment. All data are shown as mean ± SEM.

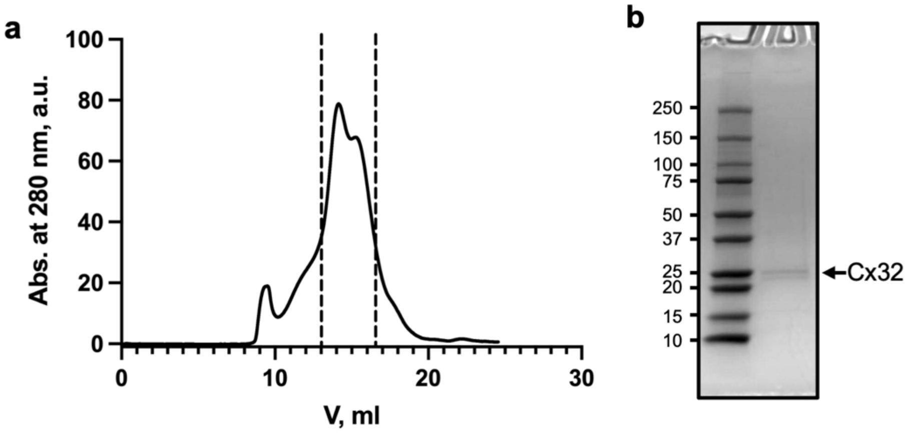


**Extended Data Fig. 3. Expression and purification of Cx32. a,** Size exclusion chromatogram (SEC) of Cx32 purification. **b,** Coomassie stained SDS PAGE gel of collected fractions from SEC.

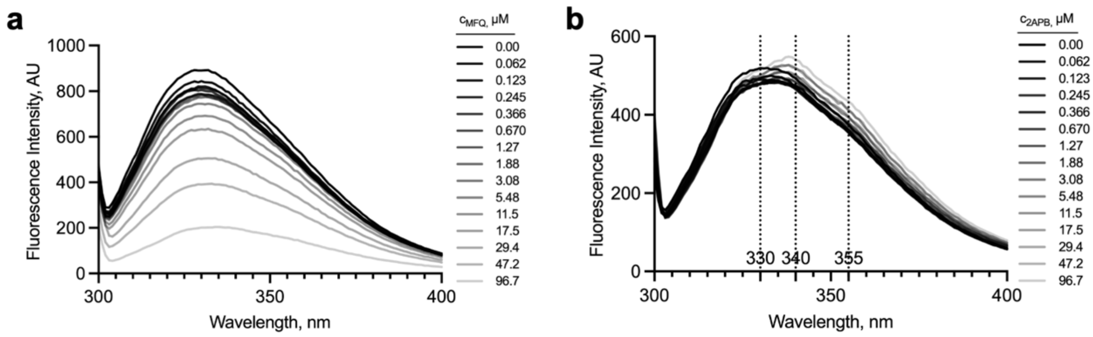


**Extended Data Fig. 4. Intrinsic tryptophan fluorescence quenching-based binding assays of MFQ and 2APB to Cx32.** Tryptophan fluorescence spectra of purified Cx32 upon titrating **a,** MFQ and **b,** 2APB (n = 3). While quenching was used for analysis of MFQ data, for 2APB the shift of the fluorescence emission curves was used to assess ligand binding.

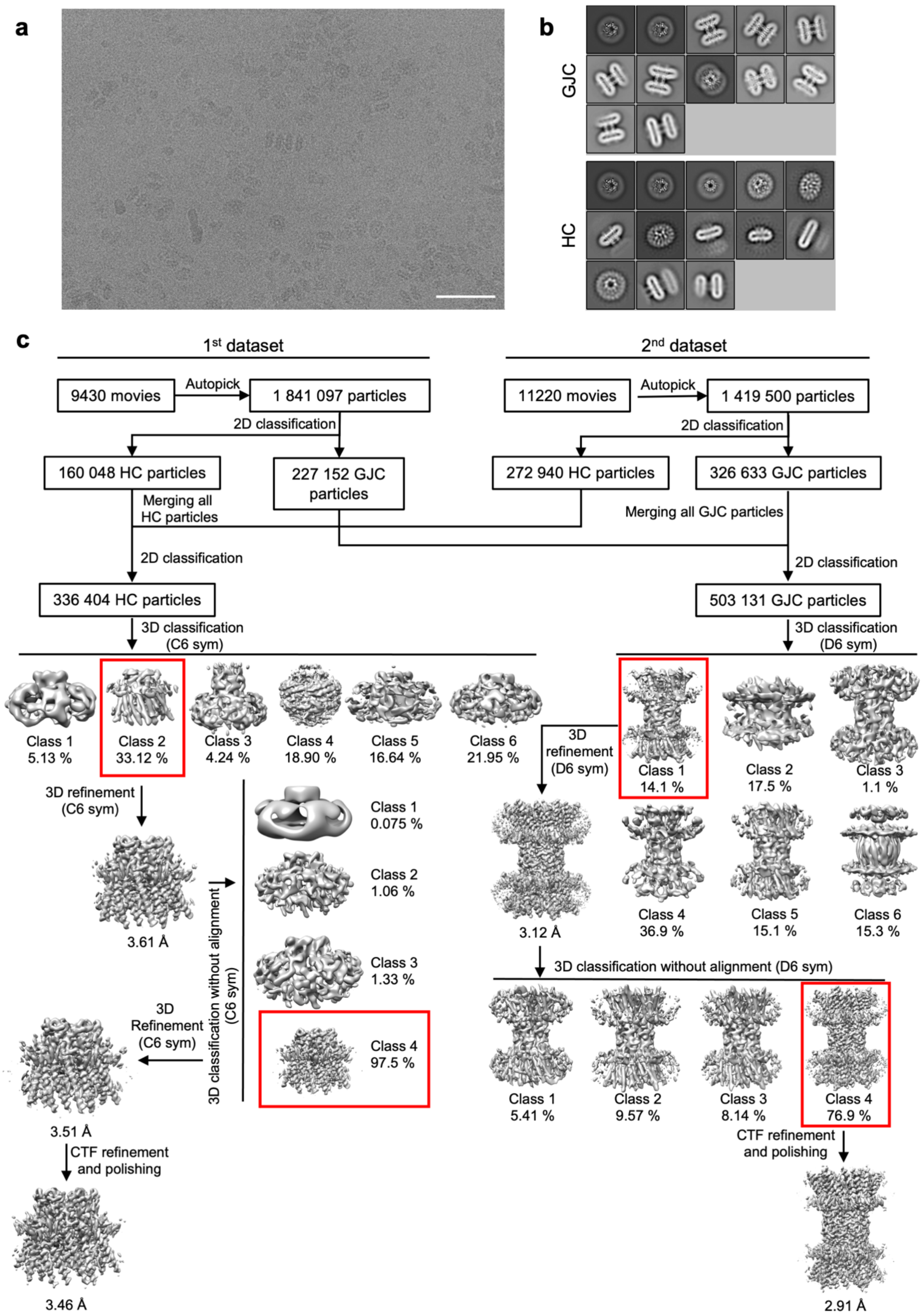


**Extended Data Fig. 5. Cryo-EM image processing pipeline of Cx32-MFQ dataset. a,** A representative cryo-EM micrograph of Cx32-MFQ dataset. Scale bar = 50 nm **b,** Selected GJC and HC classes after 2D classification. Box size = 250 Å. **c,** Overview of the cryo-EM GJC and HC processing pipeline. Selected 3D classes for further processing are indicated with red boxes.

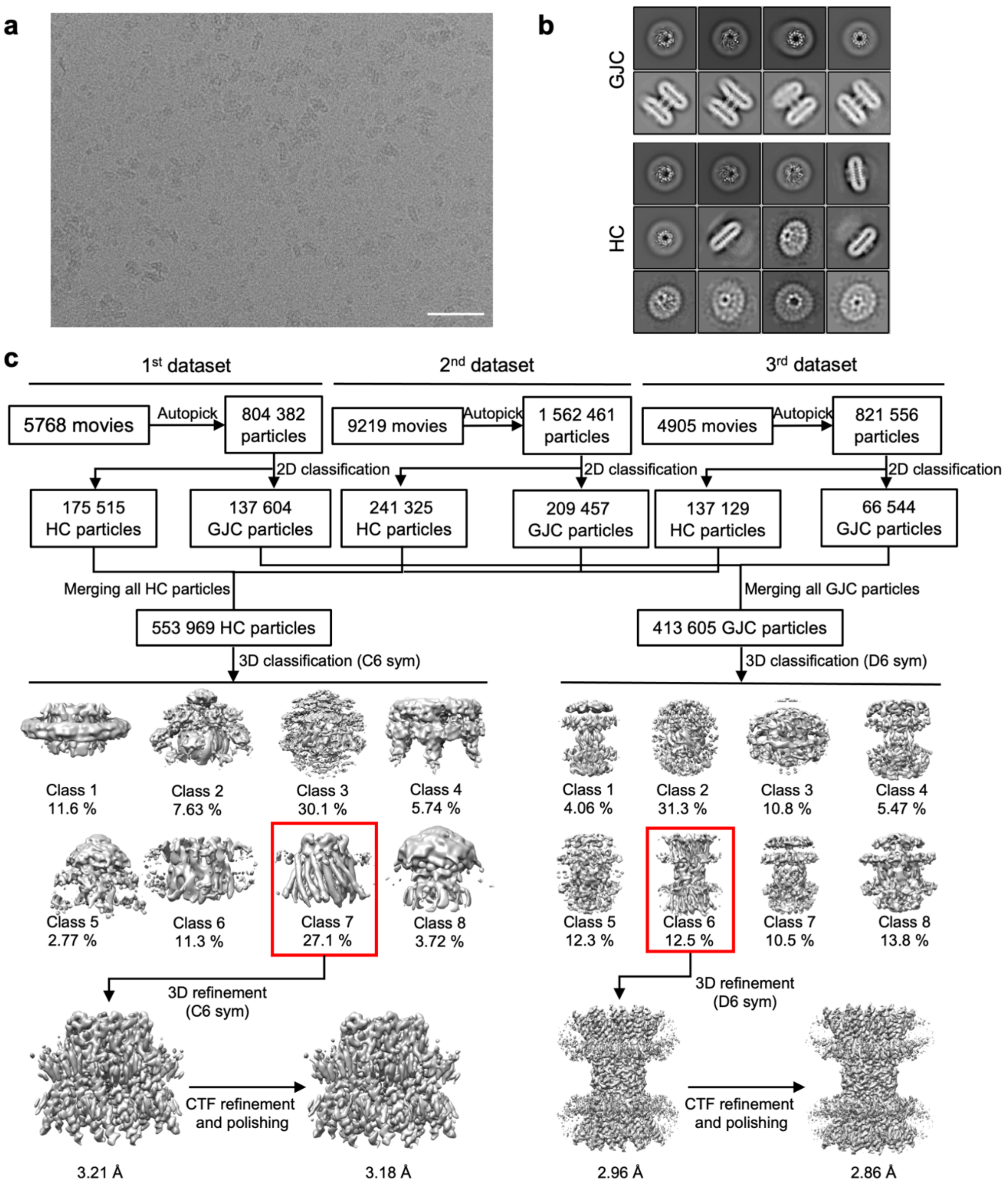


**Extended Data Fig. 6. Cryo-EM image processing pipeline of Cx32-2APB dataset. a,** A representative cryo-EM micrograph of Cx32-2APB dataset. Scale bar = 50 nm **b,** Selected GJC and HC classes after 2D classification. Box size = 250 Å. **c,** Overview of the cryo-EM GJC and HC processing pipeline. Selected 3D classes for further processing are indicated with red boxes.

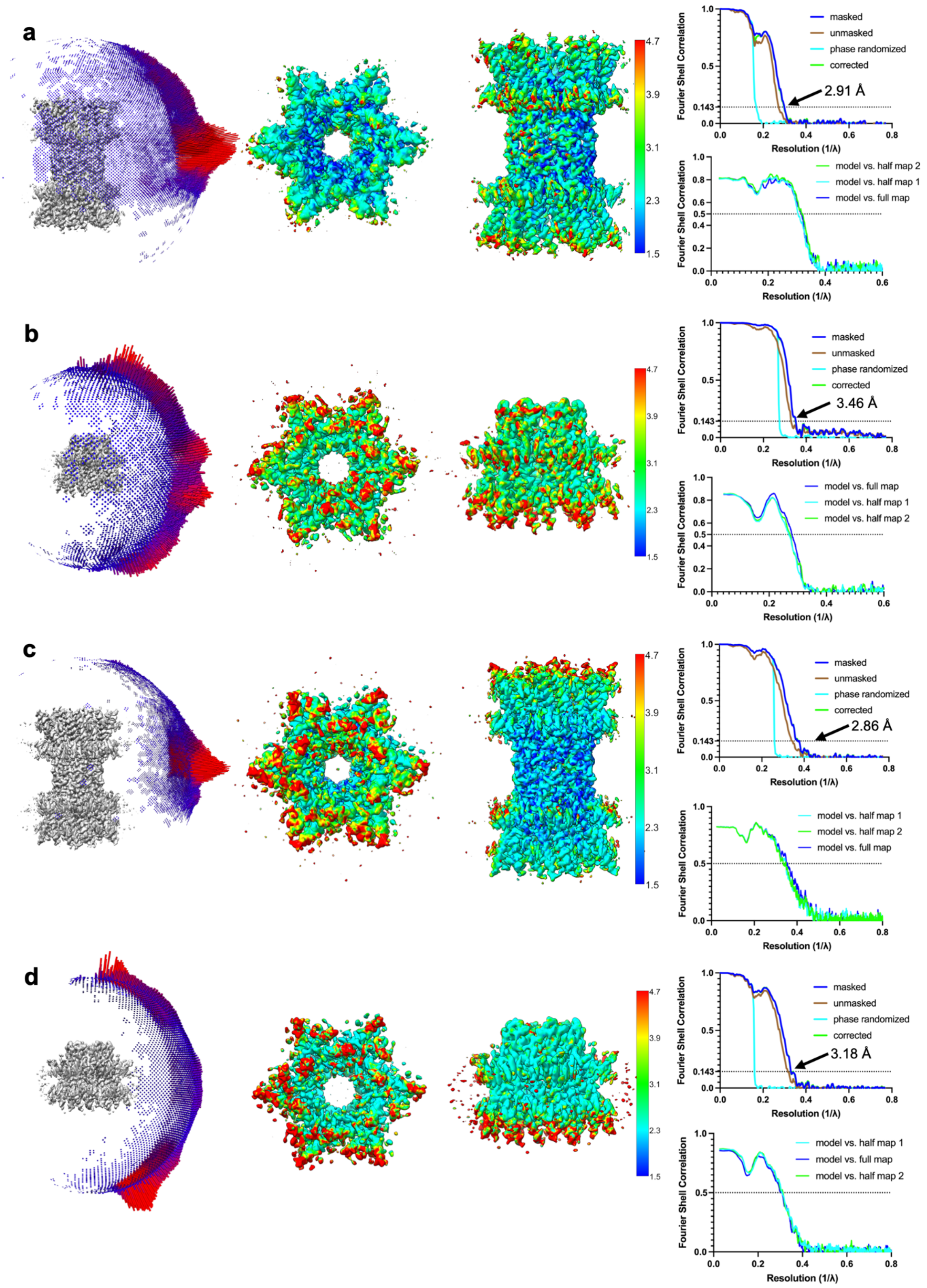


**Extended Data Fig. 7. Angular distribution, local resolution, and Fourier Shell Correlation (FSC) graphs of Cx32-MFQ and Cx32-2APB GJC and HC. a,** Cx32-MFQ GJC. **b,** Cx32-MFQ HC. **c,** Cx32-2APB GJC. **d,** Cx32-2APB HC.

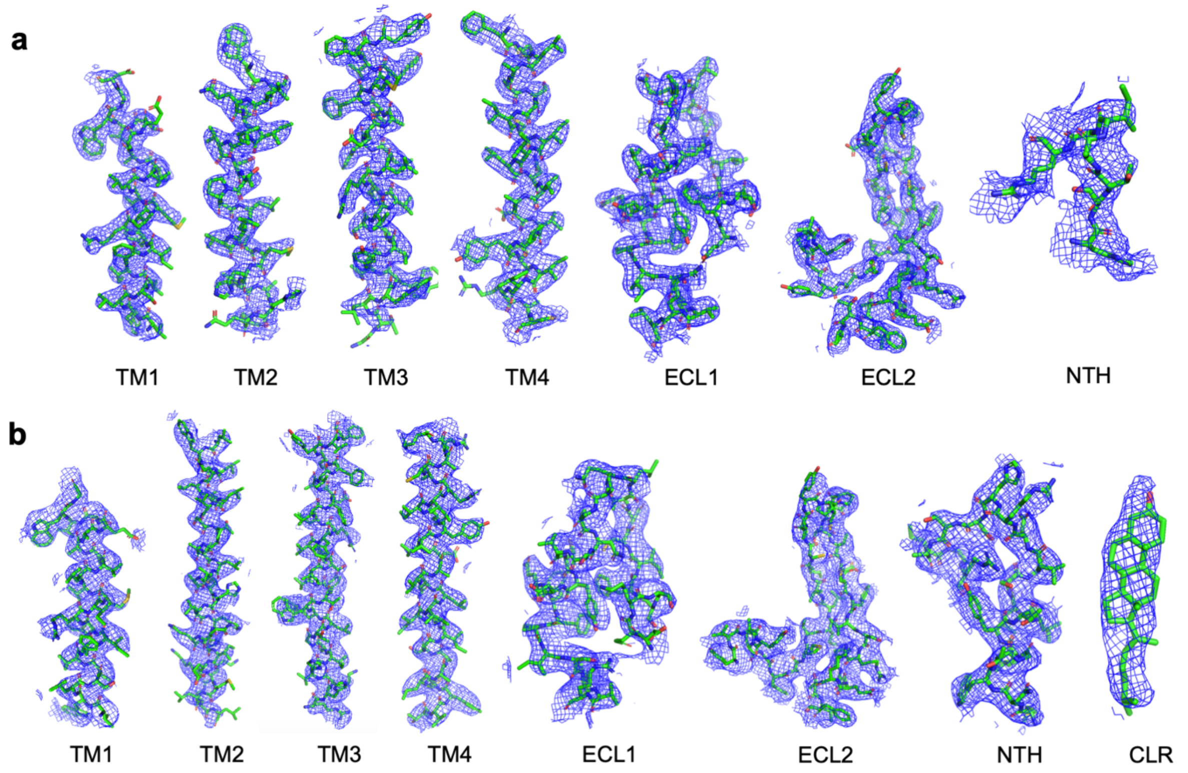


**Extended Data Fig. 8. Cryo-EM density map features of Cx32-MFQ. a,** Isolated cryo-EM density map features of Cx32 GJC in complex with MFQ, including transmembrane helices 1-4 (TM1-4), extracellular loops 1 and 2 (ECL1 and ECL2), and N-terminal helix (NTH). **b,** Isolated cryo-EM density map feature of Cx32 HC in complex with MFQ, including TM1-4, ECL1 and ECL2, NTH, and putative cholesterol (CLR) density.

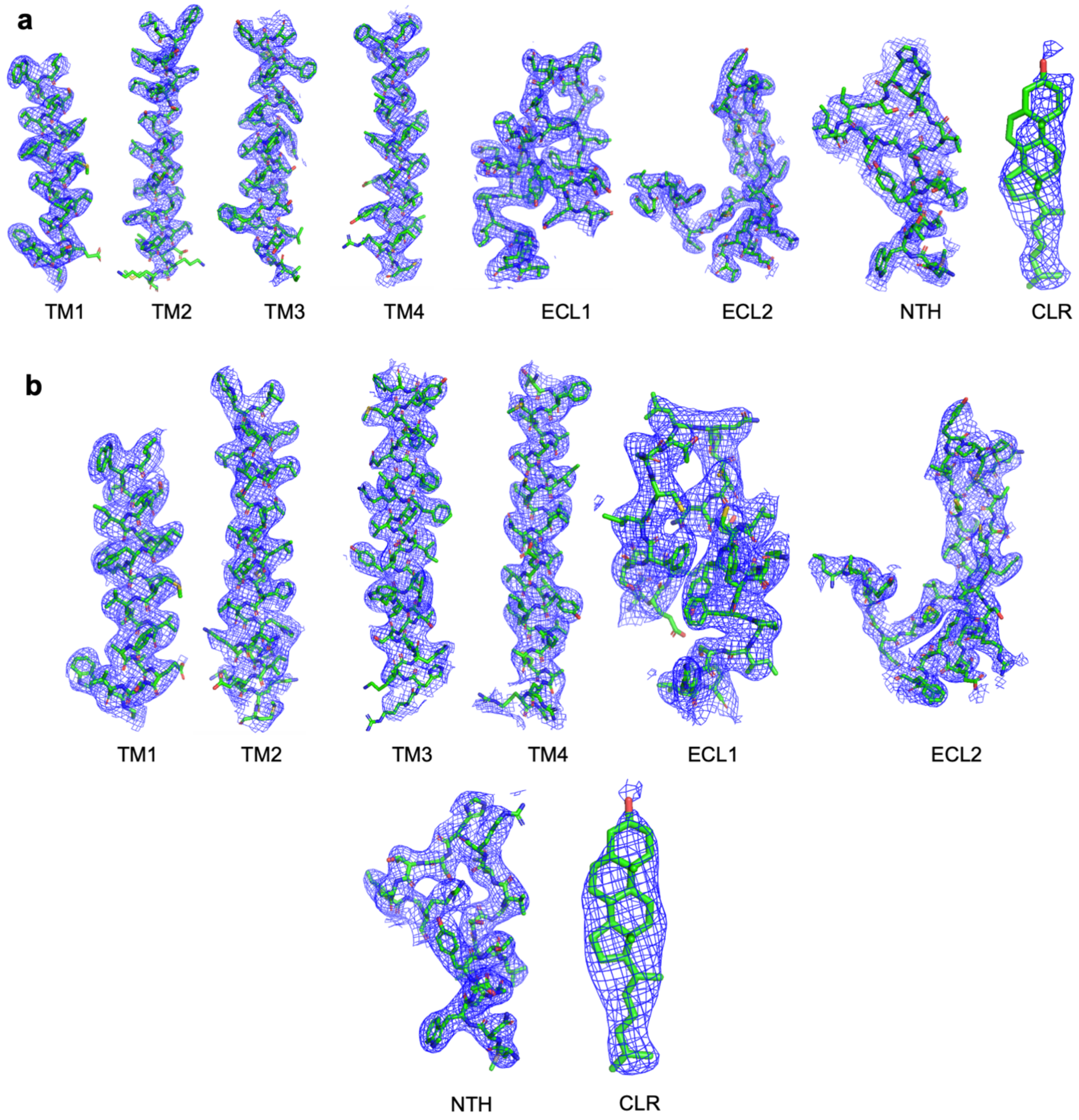


**Extended Data Fig. 9. Cryo-EM density map features of Cx32-2APB. a,** Isolated cryo-EM density map features of Cx32 GJC in complex with 2APB, including transmembrane helices 1-4 (TM1-4), extracellular loops 1 and 2 (ECL1 and ECL2), N-terminal helix (NTH), and putative cholesterol (CLR). **b,** Isolated cryo-EM density map feature of Cx32 HC in complex with 2APB, including TM1-4, ECL1 and ECL2, NTH, and putative cholesterol (CLR) density.

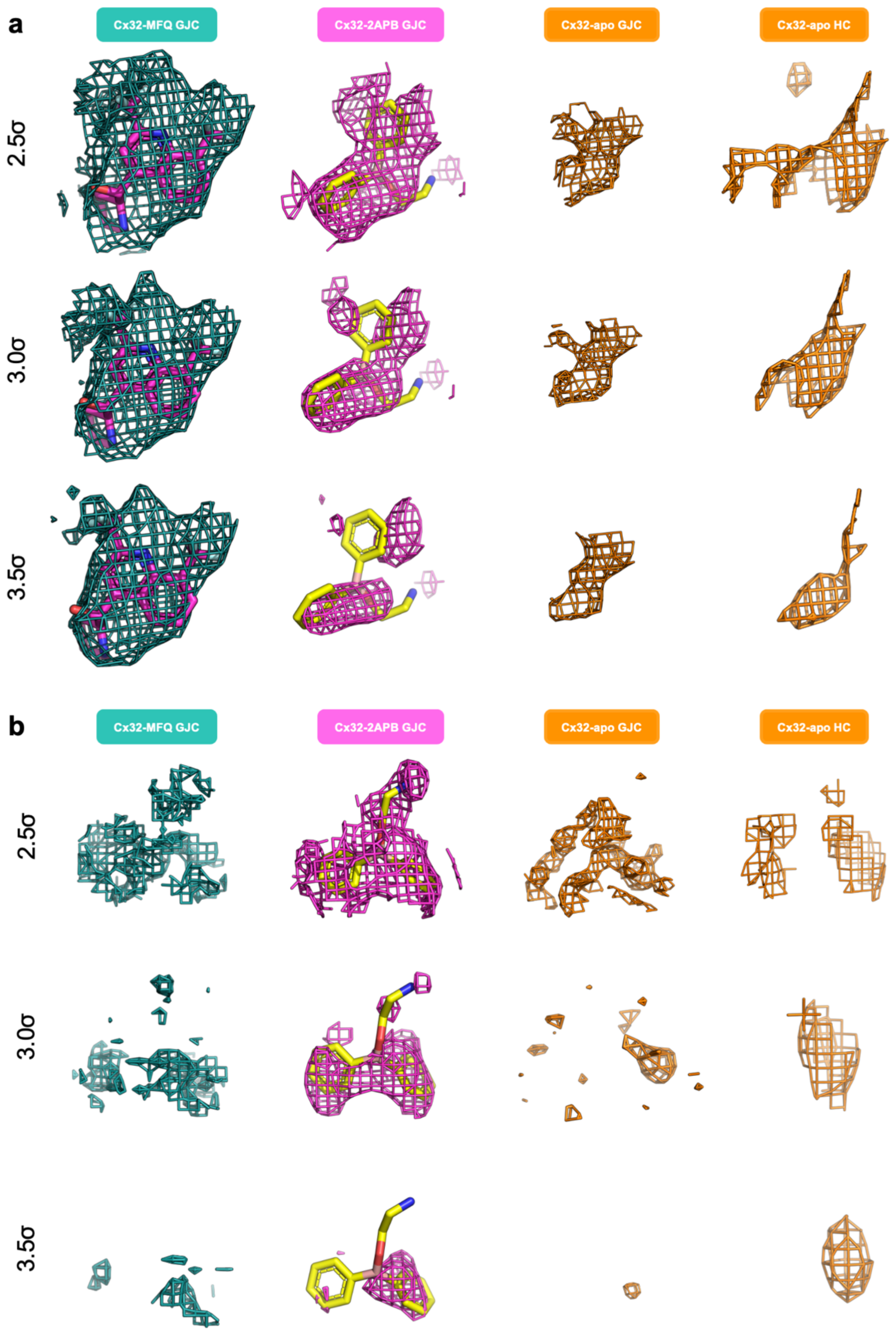


**Extended Data Fig. 10. Analysis of site A and site M densities at different contour levels.** **a,** Site M densities, contoured at 2.5σ, 3σ, and 3.5σ for Cx32-MFQ GJC, Cx32-2APB GJC, Cx32-apo GJC, and Cx32-apo HC. **b,** Site A densities, contoured at 2.5σ, 3σ, and 3.5σ for Cx32-MFQ GJC, Cx32-2APB GJC, Cx32-apo GJC, and Cx32-apo HC.

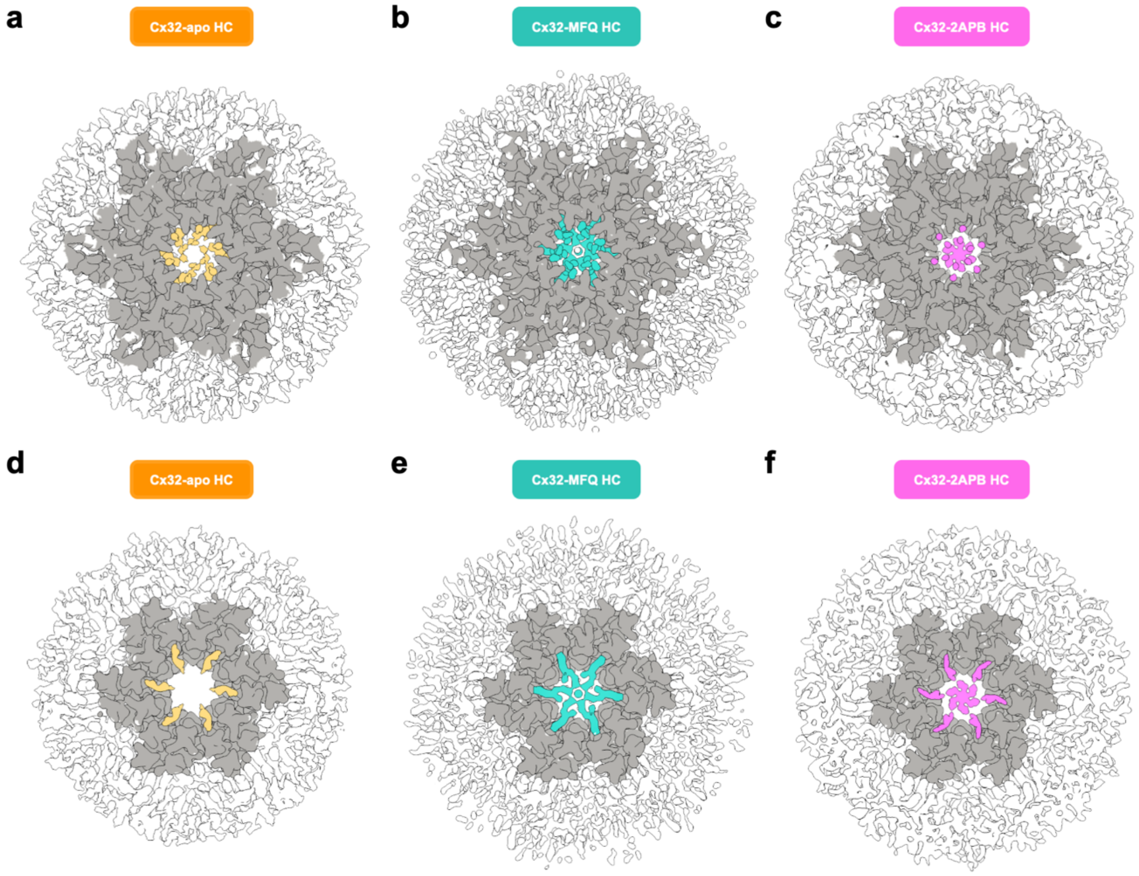


**Extended Data Fig. 11. Structural comparison of Cx32 HC in complex with MFQ or 2APB**. Top views of **a,** Cx32-apo HC, **b,** Cx32-MFQ HC, and **c,** Cx32-2APB HC density maps. Top views clipped below the NTH of **d,** Cx32-apo HC, **e,** Cx32-MFQ HC, and **f,** Cx32-2APB HC density maps. Contoured at 3σ.

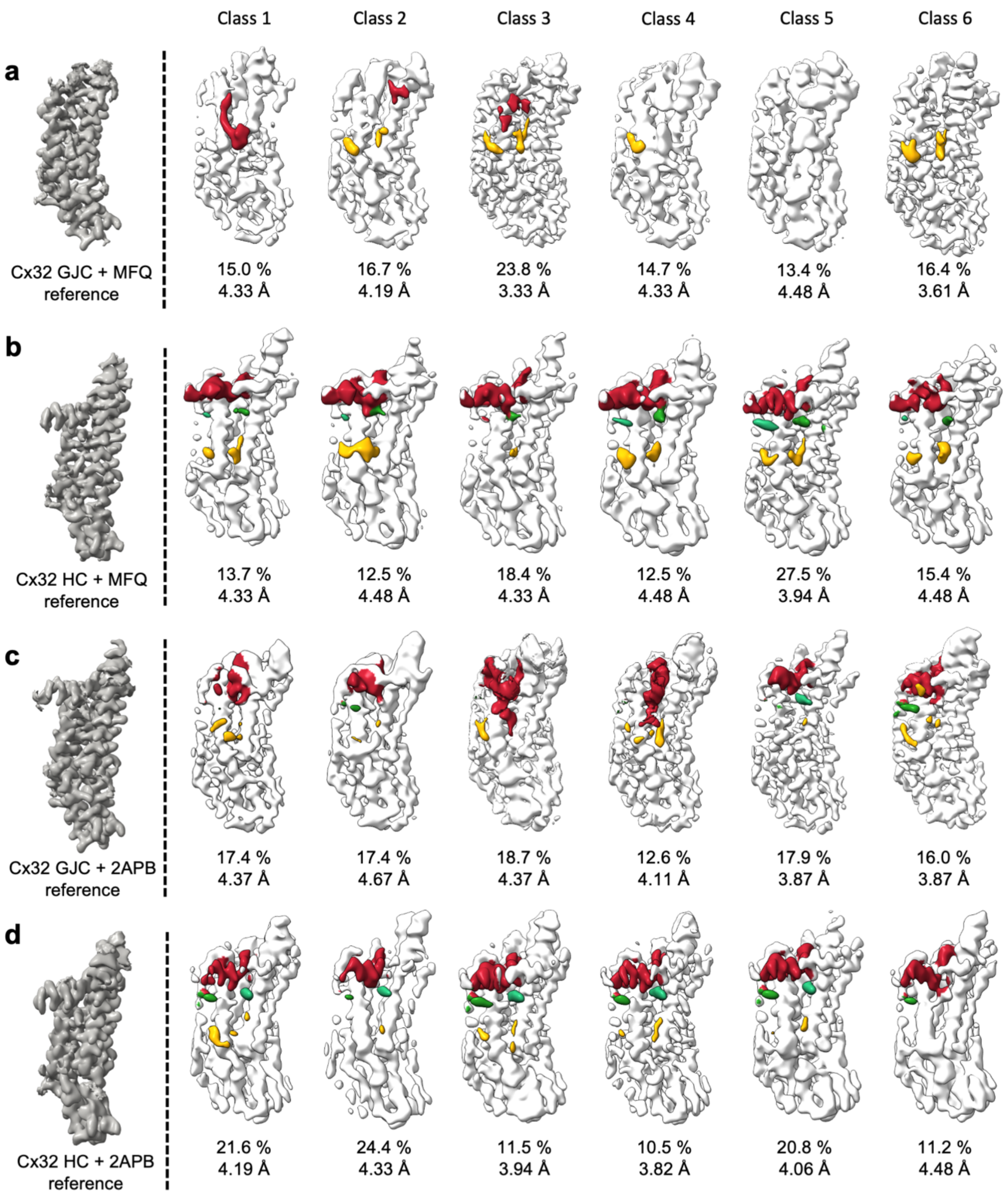


**Extended Data Fig. 12. Particle focused classification (PFC).** The NTH is depicted in red, lipid-2 in green and densities at site M in yellow. **a,** Cx32-MFQ GJC. **b,** Cx32-MFQ HC. **c,** Cx32-2APB GJC. **d,** Cx32-2APB HC.

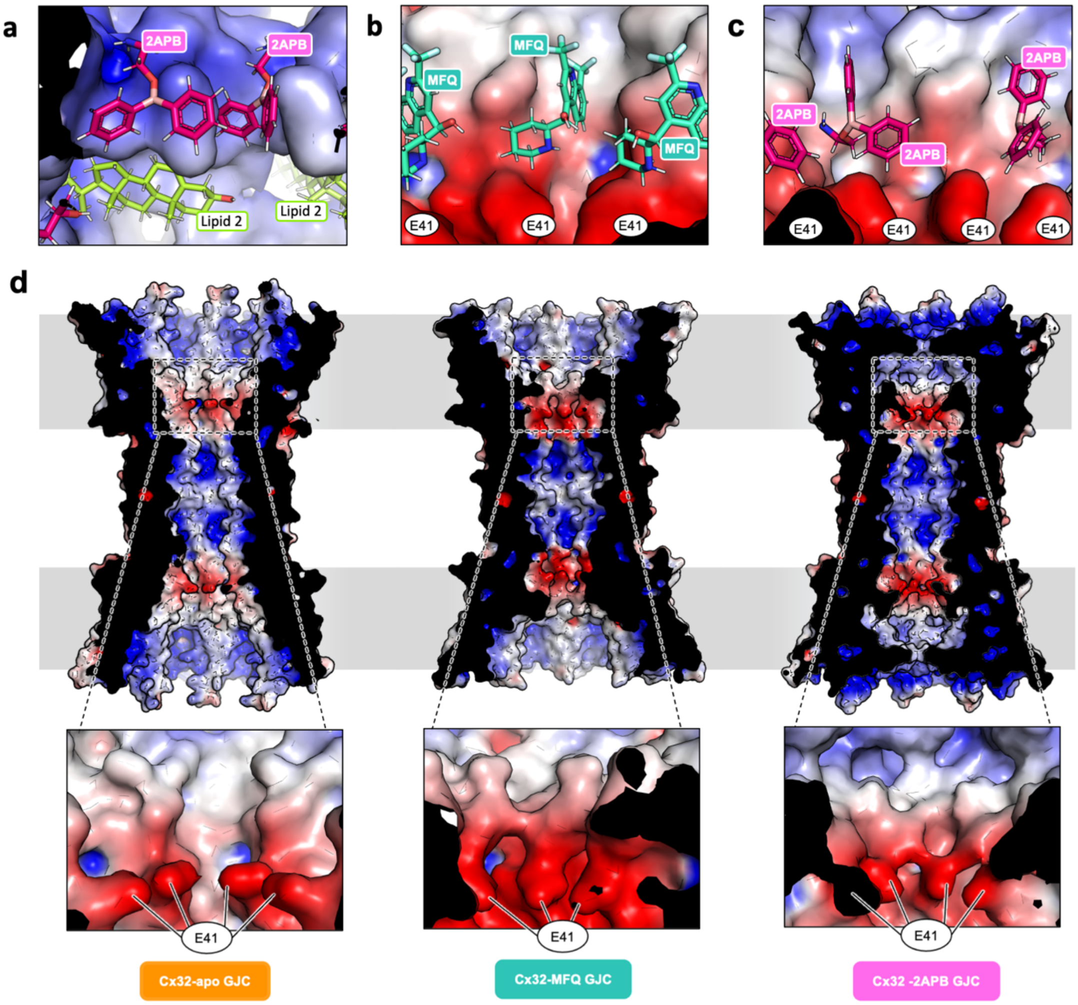


**Extended Data Fig. 13. Electrostatic surface potential analysis of Cx32 in the absence and in the presence of inhibitors. a,** Binding of 2APB and lipid-2 to site A, composed of neighboring amphipathic NTH. Binding of **b,** MFQ and **c,** 2APB to the hydrophobic site M binding pocket, positioned above residue E41. **d,** The side view of the electrostatic surface potential representation of the Cx32 without a bound drug, in complex with MFQ, and in complex with 2APB.

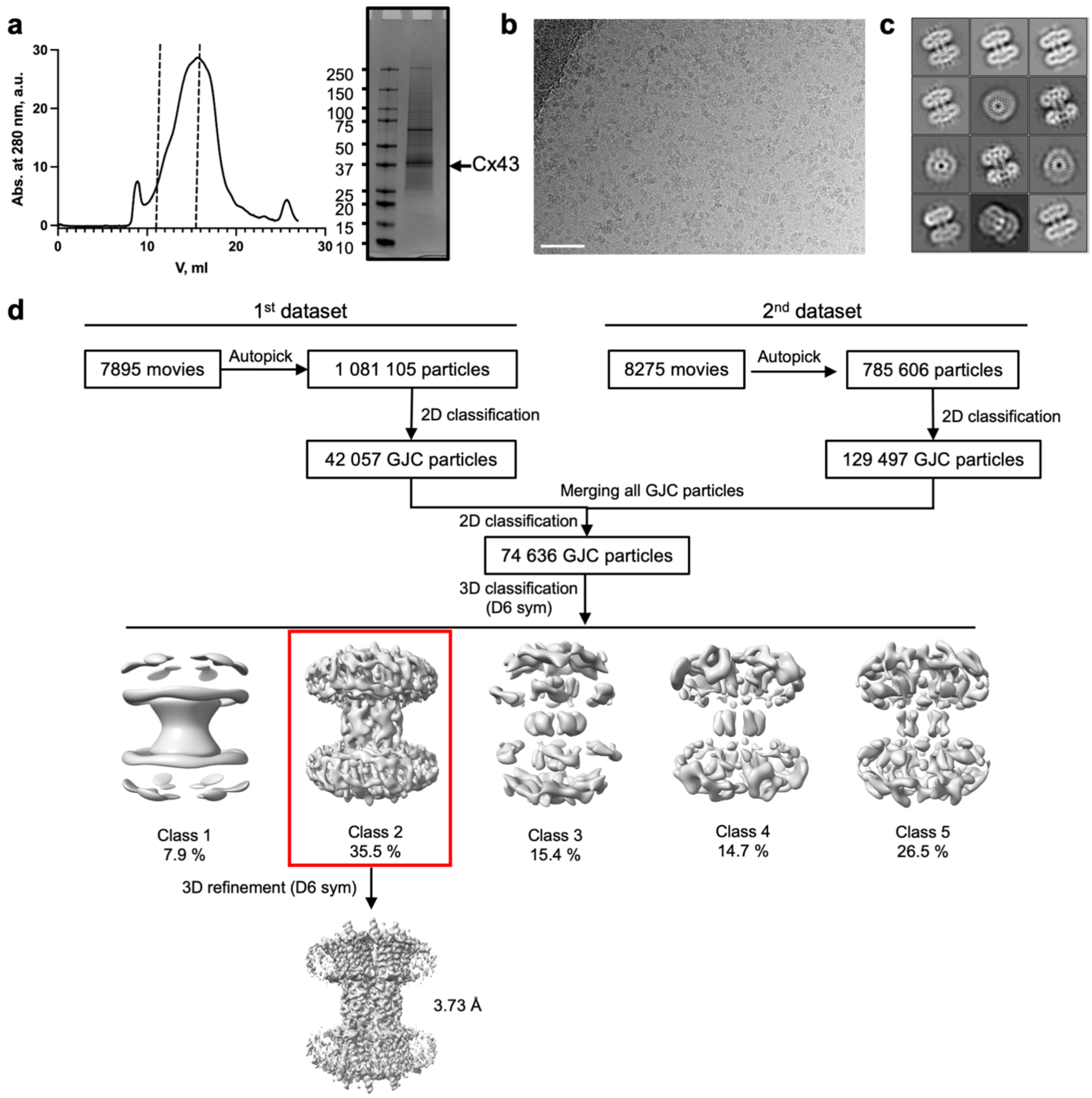


**Extended Data Fig. 14. Cx43 GJC purification and data processing pipeline of Cx43 GJC in complex with MFQ. a,** SEC profile and SDS PAGE analysis of Cx43 GJC purification. **b,** A representative cryo-EM micrograph of Cx43 in complex with MFQ. **c,** 2D classes of Cx43 GJC. **d,** Cryo-EM data processing pipeline of Cx43 GJC in complex with MFQ. Chosen classes for further processing are outlined with red.

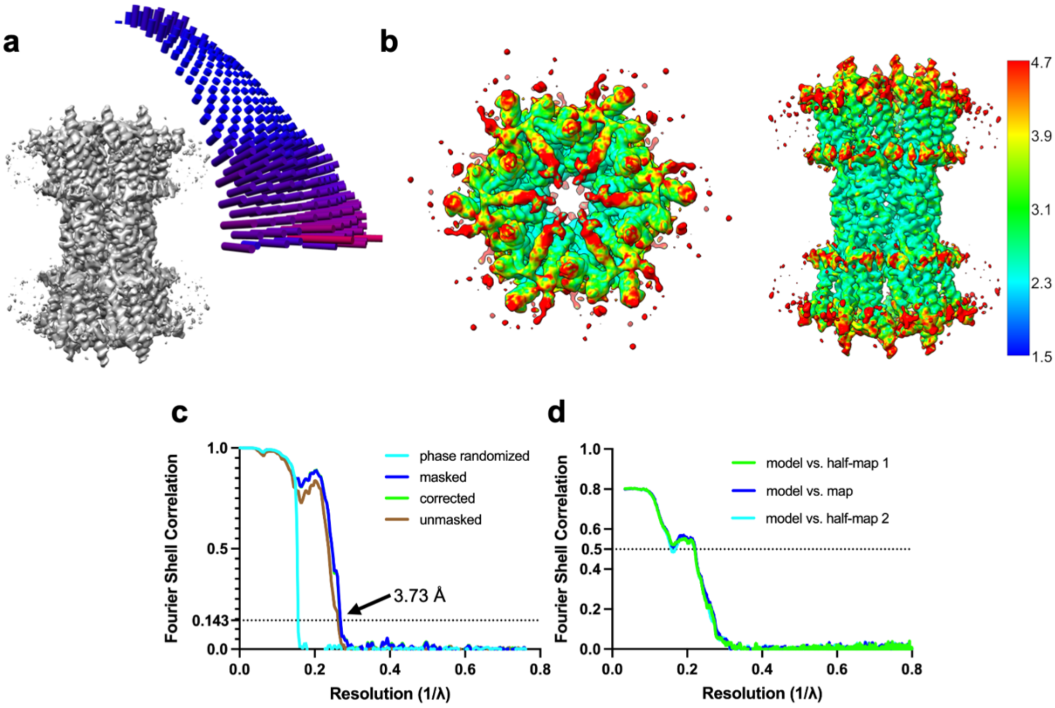


**Extended Data Fig. 15. Angular distribution, local resolution, Fourier Shell Correlation (FSC) graphs and effect of symmetry of Cx43 GJC in complex with MFQ. a,** Angular distribution. **b,** Local resolution. **c,** Fourier Shell Correlation graphs after refinement and post-processing. **d,** Fourier Shell Correlation graphs of model vs. refinement maps. **e,** Refinements of Cx43 GJC by imposing D6 symmetry.

s

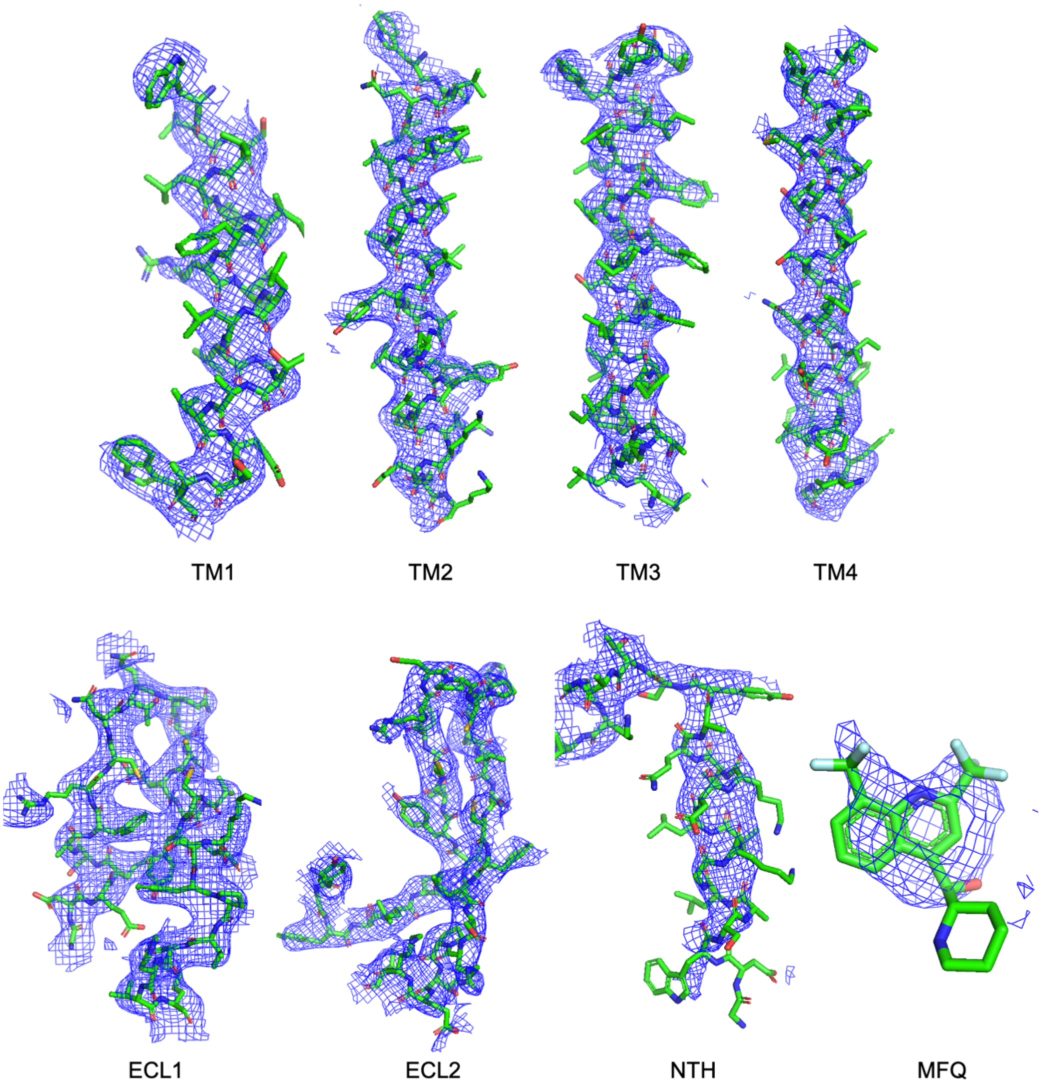


**Extended Data Fig. 16. Cryo-EM density map features of Cx43-MFQ GJC.** Isolated cryo-EM density map features of Cx43 GJC in complex with MFQ, including transmembrane helices 1-4 (TM1-4), extracellular loops 1 and 2 (ECL1 and ECL2), N-terminal helix (NTH) and mefloquine (MFQ). Contoured at 3σ.

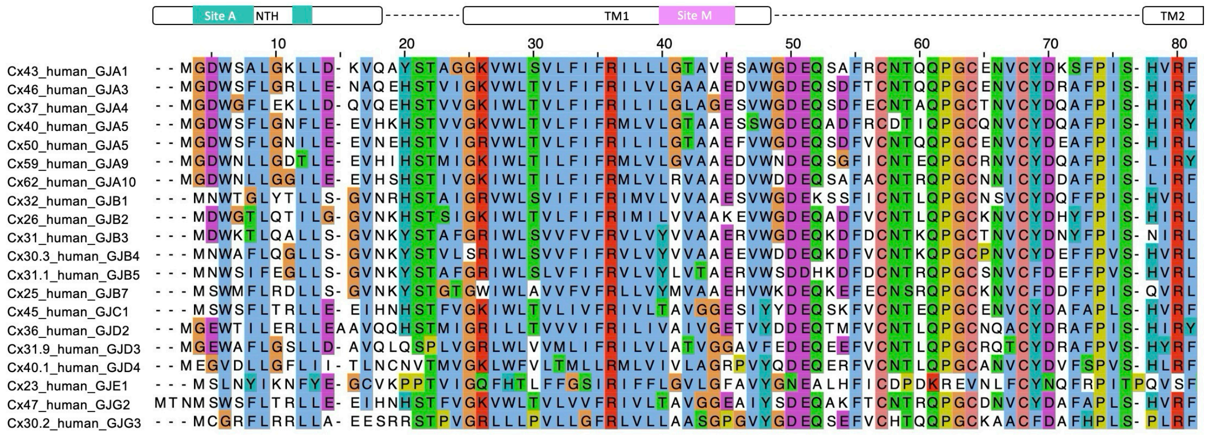


**Extended Data Fig. 17. Multiple Sequence Alignment of NTH and Site M in Connexins.** 20 sequences from connexin family are aligned and colored with Clustal scheme in Jalview. Blue, hydrophobic; red, positive charge; magenta, negative charge; green, polar; pink, cysteines; orange, glycines; yellow, prolines; cyan, aromatic; white, unconserved.

**Extended Data Table 1.** Cryo-EM data collection and processing statistics.

<table>
<tr><td colspan="8">Data collection</td></tr>
<tr><td>Sample</td><td colspan="2">Cx32 + MFQ</td><td colspan="3">Cx32 + 2APB</td><td colspan="2">Cx43 + MFQ</td></tr>
<tr><td>Instrument</td><td colspan="7">FEI Titan Krios/Gatan K3 Summit/Quantum GIF</td></tr>
<tr><td>Voltage [kV]</td><td colspan="7">300</td></tr>
<tr><td>Electron dose [e-/Å]</td><td>Dataset 1: 50</td><td>Dataset 2: 48</td><td>Dataset 1: 52</td><td>Dataset 2: 44</td><td>Dataset 3: 48</td><td>Dataset 1: 60</td><td>Dataset 2: 60</td></tr>
<tr><td>Defocus range [µm]</td><td colspan="7">0.5 to 2.5</td></tr>
<tr><td>Pixel size [Å]</td><td colspan="7">0.65</td></tr>
<tr><td>Map resolution [Å]</td><td>Cx32 GJC + MFQ</td><td>Cx32 HC + MFQ</td><td>Cx32 GJC + 2APB</td><td colspan="2">Cx32 HC + 2APB</td><td colspan="2">Cx43 GJC + MFQ</td></tr>
<tr><td>FSC threshold 0.143</td><td>2.91</td><td>3.46</td><td>2.86</td><td colspan="2">3.18</td><td colspan="2">3.73</td></tr>
<tr><td>Number of particles</td><td>56,469</td><td>108,632</td><td>51,701</td><td colspan="2">150,125</td><td colspan="2">28,940</td></tr>
<tr><td colspan="8">Refinement</td></tr>
<tr><td>Model resolution [Å]</td><td rowspan="2">3</td><td rowspan="2">3.7</td><td rowspan="2">2.9</td><td colspan="2" rowspan="2">3</td><td colspan="2" rowspan="2">4.2</td></tr>
<tr><td>FSC threshold 0.5</td></tr>
<tr><td>Map sharpening B-factor [Å]</td><td>-16.65</td><td>-126.67</td><td>-80.34</td><td colspan="2">-114.23</td><td colspan="2">-182.83</td></tr>
<tr><td>Map CC</td><td>0.88</td><td>0.89</td><td>0.94</td><td colspan="2">0.95</td><td colspan="2">0.9</td></tr>
<tr><td colspan="8">Model composition</td></tr>
<tr><td>Protein residues/ligand</td><td>2076</td><td>1242/12</td><td>2352/12</td><td colspan="2">1230/6</td><td colspan="2">2268/12</td></tr>
<tr><td>ADP (B factor)</td><td>92.85</td><td>67.03</td><td>135.61</td><td colspan="2">95.12</td><td colspan="2">170.82</td></tr>
<tr><td>Bond length r.m.s.d. [Å]</td><td>0.003</td><td>0.003</td><td>0.004</td><td colspan="2">0.004</td><td colspan="2">0.003</td></tr>
<tr><td colspan="8">Validation</td></tr>
<tr><td>MolProbity score</td><td>2.05</td><td>1.66</td><td>1.16</td><td colspan="2">1.24</td><td colspan="2">1.92</td></tr>
<tr><td>Clash score</td><td>9.65</td><td>8.1</td><td>3.68</td><td colspan="2">4.31</td><td colspan="2">9.68</td></tr>
<tr><td>Rotamer outliers (%)</td><td>5.16</td><td>0.54</td><td>0.57</td><td colspan="2">0.51</td><td colspan="2">3.14</td></tr>
<tr><td colspan="8">Ramachandran plot</td></tr>
<tr><td>Favored (%)</td><td>98.22</td><td>96.55</td><td>98.44</td><td colspan="2">98.51</td><td colspan="2">97.84</td></tr>
<tr><td>Allowed (%)</td><td>1.78</td><td>3.45</td><td>1.56</td><td colspan="2">1.49</td><td colspan="2">2.16</td></tr>
<tr><td>Disallowed (%)</td><td>0</td><td>0</td><td>0</td><td colspan="2">0</td><td colspan="2">0</td></tr>
</table>
